# ShapeRotator: an R tool for standardised rigid rotations of articulated Three-Dimensional structures with application for geometric morphometrics

**DOI:** 10.1101/159392

**Authors:** Marta Vidal-García, Lashi Bandara, J. Scott Keogh

## Abstract

1. The quantification of complex morphological patterns typically involves comprehensive shape and size analyses, usually obtained by gathering morphological data from all the structures that capture the phenotypic diversity of an organism or object. Articulated structures are a critical component of overall phenotypic diversity, but data gathered from these structures are difficult to incorporate in to modern analyses because of the complexities associated with jointly quantifying 3D shape in multiple structures.
2. While there are existing methods for analysing shape variation in articulated structures in Two-Dimensional (2D) space, these methods do not work in 3D, a rapidly growing area of capability and research.
3. Here we describe a simple geometric rigid rotation approach that removes the effect of random translation and rotation, enabling the morphological analysis of 3D articulated structures. Our method is based on Cartesian coordinates in 3D space so it can be applied to any morphometric problem that also uses 3D coordinates (e.g. spherical harmonics). We demonstrate the method by applying it to a landmark-based data set for analysing shape variation using geometric morphometrics.
4. We have developed an R tool (ShapeRotator) so that the method can be easily implemented in the commonly used R package *geomorph* and *MorphoJ* software. This method will be a valuable tool for 3D morphological analyses in articulated structures by allowing an exhaustive examination of shape and size diversity.

## Background

Data on shape and size variation is essential in many fields, including evolutionary biology and ecology, engineering, medical science, and anthropology (Loncaric 1998; McIntyre & Mossey 2003; Slice 2006). For most of these studies, the most widely used tools for analysing morphological variation within or between a group of organisms or objects are based on Cartesian coordinates of landmarks (Bookstein 1997).

Of the wide array of methods using Cartesian coordinates, geometric morphometrics (GM) is the most common, especially when analysing shape and size variation and covariation (Mitteroecker & Gunz 2009; Adams *et al.* 2013). The first two steps of this GM procedure consist of a landmark approach that: (1) gathers (two-or three-dimensional) coordinates of anatomically defined and homologous loci, followed by (2) a generalised Procrustes analysis (GPA) that superimposes configurations of each set of landmarks in all specimens, by removing all effects of size, translation and rotation, in order to only obtain shape information (Klingenberg 2008; Adams *et al.* 2013). Geometric morphometrics, therefore, allows accurate quantitative analyses of shape and size, in either Two-Dimensional (2D) or Three-Dimensional (3D) space.

3D morphological analyses are the most accurate, as objects and organisms exist in 3D space. The recent growth in x-ray micro CT scanning and surface scanning has seen a rapid increase in the application of 3D geometric morphometric techniques, but progress has been hampered by the lack of a simple method to incorporate data from complex articulated structures.

In evolutionary biology, identifying morphological differences among different groups or taxa is crucial in order to understand evolutionary processes and their relationship to the environment (Losos 1990; Ricklefs & Miles 1994; Pagel 1999). This can be difficult, especially if traits have co-evolved, or if morphological diversification has been hindered by phylogenetic legacy or trade-offs imposed by the organism’s functional habitat (Ghalambor *et al.* 2007). Complex body shape patterns require more detailed analyses of shape, obtained by collecting data from several structures that capture the whole gamut of morphological variation in an organism. One example of this is the extraction and assembly of data from articulated structures, such as skeletons, for 3D analyses with geometric morphometric techniques. This is especially important in functional morphological studies, as they usually involve analysing more than one structure due to mechanical correlations or morphological integration. For example, jointly analysing the skull and mandible could be crucial to disentangle the relationship between diet and head shape evolution (Cornette *et al.* 2013). Similarly, collectively evaluating different modules in the limbs, especially when correlated to locomotion, or considering several structures across the whole body, could improve our understanding of the effect of environmental conditions on morphological evolution (Vidal-Garcia & Keogh 2017).

Unfortunately, non-rigid structures, such as articulated structures, will inevitably suffer the effects of natural or free rotation or translation events and be different in each individual and structure (Adams, 1999). These events could obstruct the correct quantification of shape variation by adding rotation artifacts to GM analyses (Adams *et al.* 2004). Thus, orientation of these structures needs to be corrected and standardised prior to performing shape analyses. Several approaches for shape analysis of landmark data in articulated structures already have previously been proposed (Adams 1999; Bookstein 1991). However, even though these approaches can be used in 2D and also in 3D data with further modifications, currently it has only been implemented in the two-dimensional space, such as in *geomorph*’s function fixed.angle()(Adams & Otárola-Castillo 2013).

Here we present the R tool *ShapeRotator*: a simple geometric rigid rotation approach to study 3-Dimensional (3D) shape of articulated structures, or independent structures, within an organism. We describe a method that removes shape variation due to the effect of translation between independent structures and rotation generated by movement in an articulation, among others. Thus, our approach translates and rotates articulated (or even independent) structures in order to obtain a comparable data set with all effects of random movement and rotations removed.

We apply this method to two landmark-based data sets for analysing shape variation using geometric morphometrics: (A) two sub-structures in a single-point articulation, and (B) two sub-structures in a double-point articulation (Fig. 1, 2a,b). We also provide the example data set used in *ShapeRotator* (available in github) to execute these two kinds of rigid rotations, which then allows geometric morphometric analyses to be performed in the two most commonly used 3D GM analytical software packages: *geomorph* (Adams & Otárola-Castillo 2013), and *MorphoJ* (Klingenberg 2011). This method also will allow exporting of the rotated coordinates for posterior analyses in other software platforms, even outside of the field of geometric morphometrics. Since the basis of this method lies upon rigidly spinning any structure defined by 3D coordinates, it could be used in any other shape analyses that use coordinate data, such as continuous surface meshes used in spherical harmonics (Shen *et al.* 2009). Our method is a convenient addition to the rapidly evolving tool kit of geometric morphometrics because it allows a more comprehensive exploration of morphological diversity through the gathering of shape data from complex 3D structures.

**Fig. 1.**
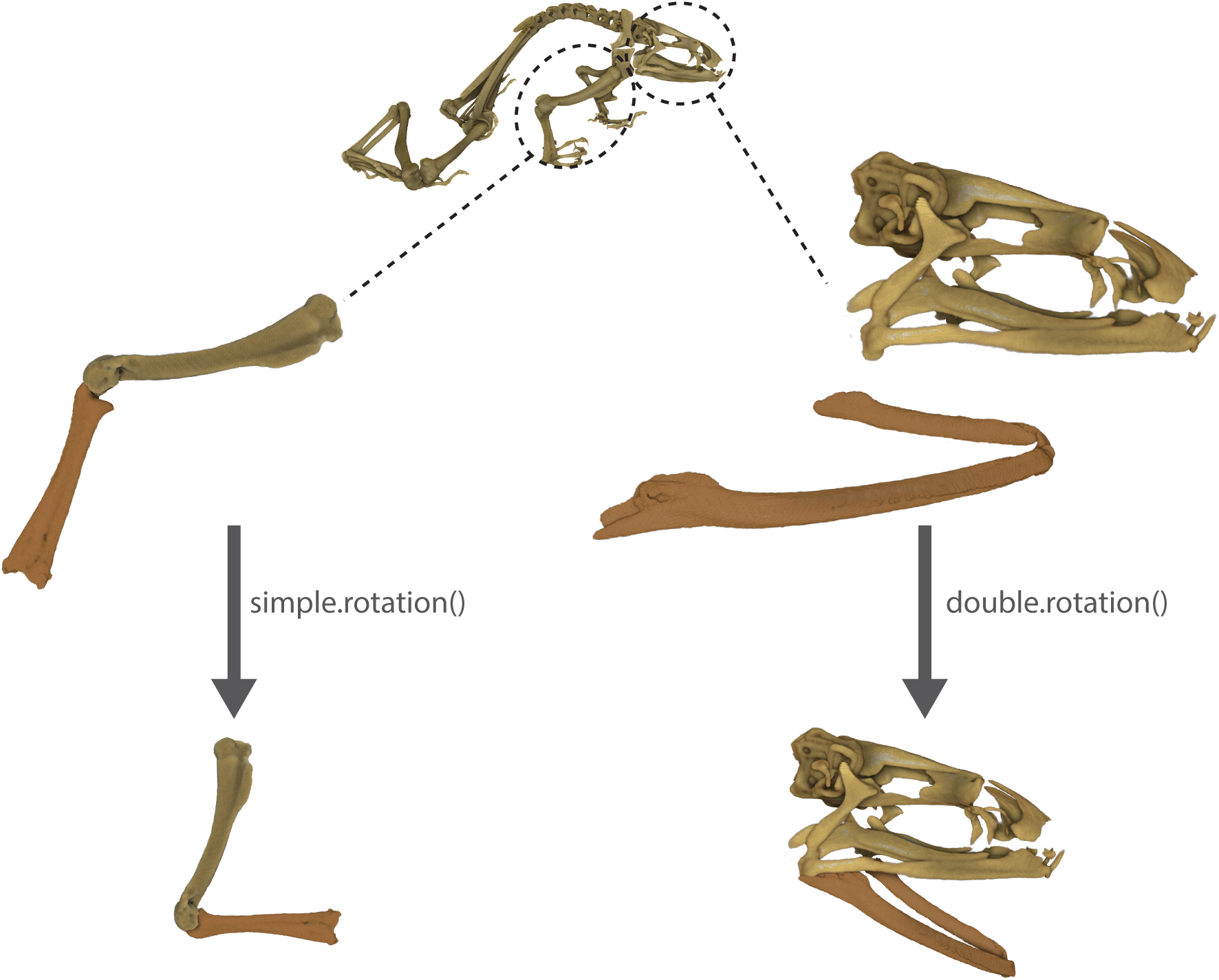
Two structures in a simple-point articulation and a double-point articulation being rotated using *simple.rotation()* and *double.rotation()*, respectively. The structures depicted are a frog humerus and radioulna for the single-point articulation and a skull and a detached mandible for the double-point articulation.

**Fig. 2.**
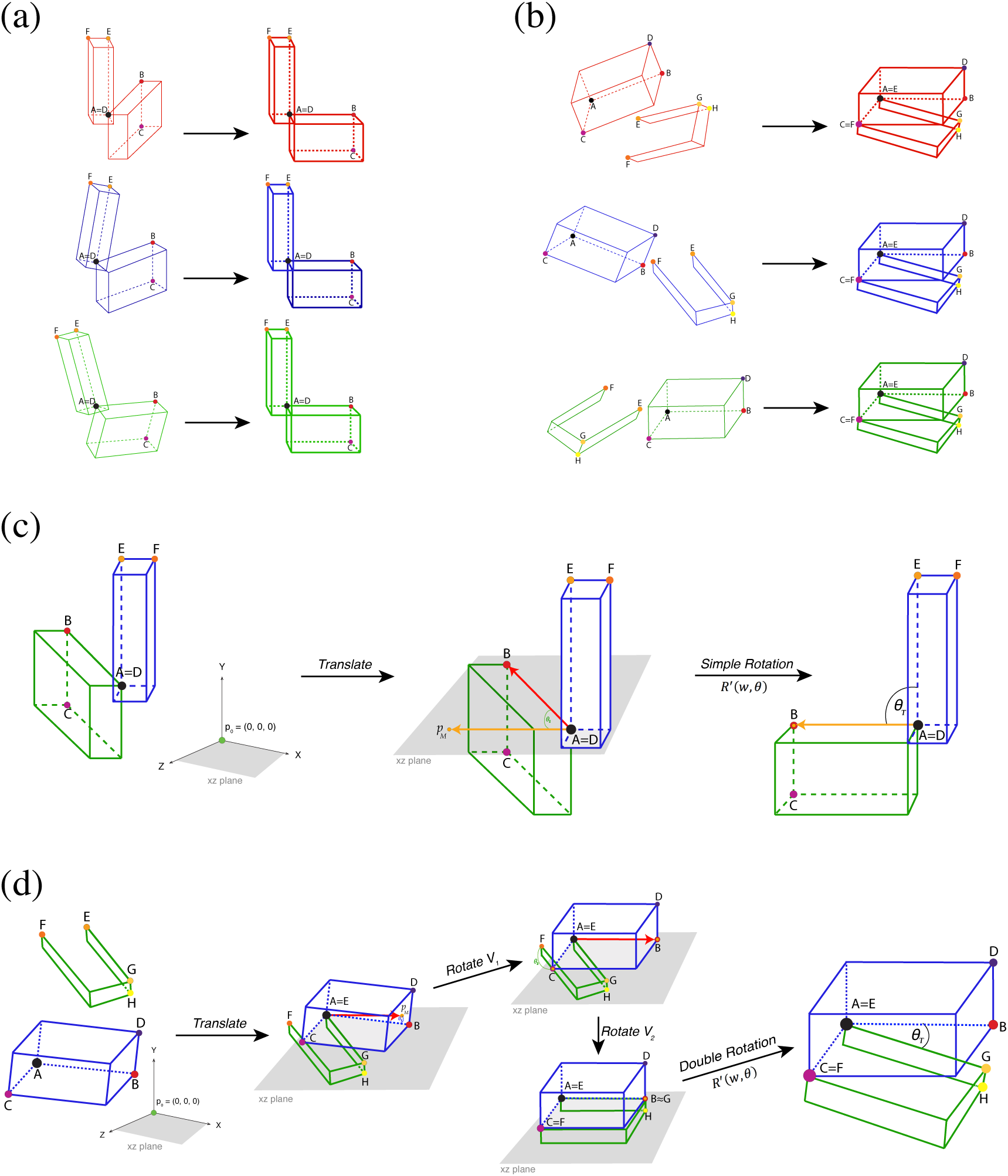
(a) Application of a translation -*translate()*-and the 3D rigid rotation method *simple.rotation()* in a simple-point articulation (e.g. humerus and radioulna) for three different ‘specimens’, by rotating articulated structures to a standardised position relative to each other. (b) Application of the translation -*translate()*-and the 3D rigid rotation method *double.rotation()* in a double-point articulation for three different ‘specimens’ (e.g. skull and mandibles). (c) Rotation method exemplified for *simple.rotation()* by depicting the plane spanned by the already translated point *p*_0_ and *A*. Please note that *p*_0_ depicts the origin point (0, 0, 0). The rotated resulting point *p*_*M*_, landmarks B, C, D, E and F, and angle θ_T_ (desired angle between the two structures) are also depicted. (d) Rotation method exemplified for *double.rotation()*. Please note that since both structures are not attached for the double-point articulation example, they are both translated to *p*_0_, which depicts the origin point (0, 0, 0). The rotated resulting point *p*_*M*_, landmarks A, B, C, D, E, F, G and H, and angles θ_1_, θ_2_ and θ_T_ (desired angle between the two structures) are also depicted.

## Methodology

We begin with a set of points 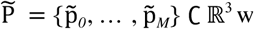 which represents a 3D object, and are ordered so that 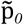 represents the base point and 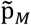 represents the end point, by which we mean that this object has an axis starting from 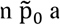 and ending at 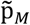. Our goal is to rotate these points via a rigid motion so that the axis on which these two points sit is either on the *X, Y* or *Z*-axis in ℝ^3^,. Rotation of vectors in ℝ^3^, is a well-known and easily resolved problem, and various formalisms exist in geometry. Thus, we translate our set of points 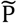 so that 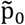 maps to the origin (0, 0, 0). This is a simple transformation *T* defined by:

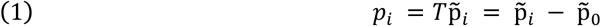

Note that the axis *X =* span {(1, 0, 0)}, *Y =* span {(0, 1, 0)}, *Z =* span {(0, 0, 1)}, where each of the generating vectors are unit. Let us fix our desired axis to which we rotate the object to be *A =* span *a* where *a =* (1, 0, 0), or *a =* (0, 1, 0), or *a =* (0, 0, 1). Since we have translated points {p_i_} and vectors correspond to positions, we are simply looking to rotate the vector *p*_*M*_ to *A*, and each other vector as a rigid motion with respect to this rotation. There are a number of ways to do this, but the simplest way is to consider the plane spanned by *p*_*M*_ and *A*, and then to rotate by the angle between *p*_*M*_ and *A* within this plane (Fig. 2c,d). Such a rotation is done via rotating on the axis to the plane, which is determined by a normal vector to this plane.

Let us describe this set-up slightly more generally. For two vectors *u, v ∈* ℝ^3^, the axis to the plane spanned by these two vectors is determined by a unit normal to the plane (there are two choices due to orientation), which we denote by *N*(*u, v*):

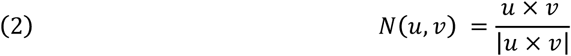

where × is the *cross product*. The angle between these vectors is then given by ∠(*u, v*):

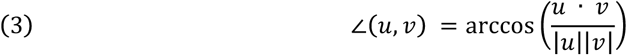

where · is the *dot (scalar) product* between vectors. The rotation matrix about an axis *w ∈* ℝ^3^, where

*w = w*_1_, *w*_2_, *w*_3_ is a unit vector, of angle θ radians is given by the well-known matrix:

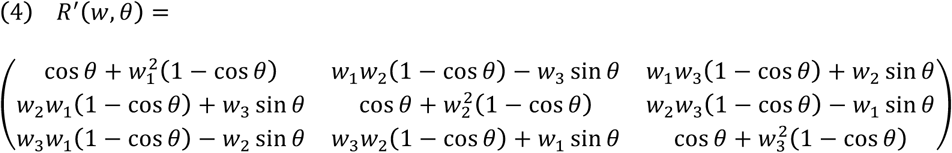

Thus, to obtain a rotation matrix which is the rigid motion rotating the vector *u* to *v* in the plane spanned by *u* and *v*, we obtain the expression:

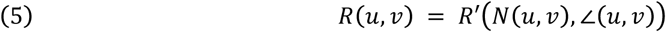

Getting back to our original problem, we set *v = p*_*M*_ and *u = a*, and then we have the rotated points:

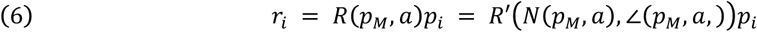

where *R* (*p*_*M*_, *a*) *p*_*i*_ is the action of the matrix *R* (*p*_*M*_, *a*) on the vector *p*_*i*_.

It may be necessary to introduce a further constraint in the rotation. For instance, suppose *a =* (0, 1, 0) and there is a point *p*_*I*_, now rotated to *r*_*I*_ via the method we describe, which should lie in the *Y*-axis. That is, we need to further rotate *r*_*I*_ to a point 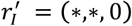. To do this, we simply rotate in the axis *a*, by an angle θ_Y_ *r*_*I*_ *=* arctan ((*r*_*I*_)_3_ / (*r*_*I*_)_1_), where *r*_*I*_ *=* ((*r*_*I*_)_1_, (*r*_*I*_)_2_, (*r*_*I*_)_3_. That is,

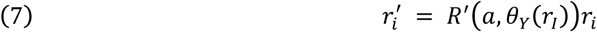

## Implementation

### Overview of the *ShapeRotator* package

Here we illustrate the functions available within the *ShapeRotator* R tool and the basic steps required in order to successfully implement the rotation on a data set of 3D coordinates. *ShapeRotator* allows the rigid rotation of sets of both landmarks and semi-landmarks used in geometric morphometric analyses, enabling morphometric analyses of complex objects, articulated structures, or multiple parts within an object or specimen. This tutorial uses two example data sets: (A) two neighbouring bones of the arm in a frog (humerus and radioulna), representing a single-point articulation example, and (B) a skull and a mandible, representing a two-point articulation example (Fig. 1). The main steps required are: (1) importing the data and fixating the rotation axes, (2) translating the whole data set of coordinates or points so that the main selected point 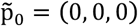, and (3) rotating the two structures to the desired angle (as outlined on Fig. 2a,b).

### Importing a data set

In the first example data set (example A) we use two geometric morphometric data sets containing both landmarks and semi-landmarks for two neighbouring and articulated bones (humerus and radioulna) from a group of several species of frogs (details in Appendix S1), in tps format. We first import the data sets using the R package *geomorph* (Adams & Otárola-Castillo 2013):

library(devtools)

install_github(“marta-vidalgarcia/ShapeRotator”) library(geomorph)

radioulna <-readland.tps(“radioulna.tps”, specID = “ID”, readcurves = F)

humerus <-readland.tps(“humerus.tps”, specID = “ID”, readcurves = F)

These two GM data sets (radioulna and humerus) will be rotated on different rotation axes in order to conform the aimed angle between them. This process is not exclusive to two neighbouring structures, and thus, it could be performed for as many independent subunits as desired by choosing the different angles among different rotation axes and all the translation processes. For more help on importing the GM data sets please see Adams *et al.* (2014), and the associated help files. Please note that this method also works for semi-landmarks as long as they have been equidistantly positioned prior to the translation and the rigid rotation. The same process is needed for the second example dataset (example B).

### Translating

During this step each structure will be translated to the point of origin so that 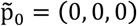, thus the distance from the coordinates of landmark_a *A*_*x*_, *A*_*y*_, *A*_*z*_ is substracted from all the landmarks in all specimens, e.g. *N*_*x*_ - *A*_*x*_, *N*_*y*_ - *A*_*y*_, *N*_*z*_ - *A*_*z*_ for landmark N. This translation is made with the function *translate()*, as it follows for the one point articulation example (example A):

> data.1_t <-translate (T=data_1, landmark=landmarkA)
>
> data.2_t <-translate (T=data_2, landmark=landmarkD)

And for the double-point articulation (example B):

> data.3_t <-translate (T=data_3, landmark=landmarkA)
>
> data.4_t <-translate (T=data_4, landmark=landmarkE)

Please note that the default is to set the origin point to 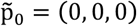, but this can be changed to another origin point. For example:

> data.1_t <-translate (T=data_1, landmark=landmarkA, origin = c(1,3,5))
>
> data.2_t <-translate (T=data_2, landmark=landmarkD, origin = c(1,3,5))

### Fixing the rotation axis

In order to fix a rotation axis in a structure we first need to select in our data set two suitable landmarks for each structure which the rotation axis will go through: landmarks A and B (for the first structure), and landmarks D and E (for the other structure). In the radioulna data set from example A, landmark A is the landmark in the 1^st^ position and landmark B is in the 10^th^ position. Similarly, for the humerus data set, landmark D would be the landmark on the 52^nd^ position, and landmark E would be in the 19^th^ position. Please note that in example A, we would perform a simple rotation between two structures, based on the angle formed by three points (one on each structure and the articulation point itself. However, since we aim to remove all articulation-related variation, including torsional rotations and mirroring artefacts, we need an extra landmark in each structure that shares at least one of their coordinates (but ideally two) with the distal landmark (e.g. y_A_ = y_B_ for the first structure, and z_D_ = z_E_ for the second). This extra landmark is needed for the simple reason that there is not enough information about the orientation of the structure with only two landmarks per structure, so even though the rigid rotation will work properly, it could position this structure in the wrong ‘mirroring’ orientation (Fig. 2c). This orientation issue is corrected by adding landmark C. In this example, landmark C is the 17^th^ landmark, while landmark F is the 107^th^ landmark in the humerus data set. We need to know which landmarks will be selected in both structures prior to the rotation process in order to ensure that the rotation process will work properly. Setting up landmarks that will be used in the rotation process of Example B is very similar to Example A, but it needs four landmarks for each structure instead of three, as it is a more complex articulation that requires two separate rotations per structure (Fig. 2d). Even though each of these rotations are calculated internally (only the four landmarks per structure and the desired angle between them need to be provided), it will be beneficial to choose landmarks that are spatially arranged in a way that facilitates the rotation process, and results in the rotating multi-structure being placed in a biologically-relevant angle between each sub-structure. ShapeRotator will give a warning message if the landmarks chosen are not optimal.

### Rotating

In the rotation step, we will use the function *simple.rotation*() in order to rigidly rotate the two structures of example A to the desired angle (in degrees), as it follows:

joined_dataset <-simple.rotation(data.1 = data.1_t, data.2 = data.2_t, land.a = 10, land.b=1, land.c=17, land.d=52, land.e=19, land.f=107, angle = 90)

The input datasets data.1 and data.2 correspond to the two translated datasets (in this case translated_radioulna and translated_humerus. We then use the selected landmarks as explained in the previous section. Finally, we include the angle (in degrees) that we would like to use to position the two structures to one another. One of the options of the function *simple.rotation*() is to select the desired angle between the two structures so that we can perform the rigid rotation of each structure positioning them in the selected angle in relation to each other. In order to do so we use the internal function *vector.angle()*, by providing the desired angle in degrees (from 0° to 360°. The function *vector.angle()* will return angle.v, a vector that forms that angle with the vector (1, 0, 0). In the example data set in ShapeRotator we rotate the coordinates from the two bones so that they form an angle of 90° degrees to each other:

New_vector <-vector.angle(90)

So that New_vector = c(0, 1, 0). Thus we could check the vector that the function *simple.rotation*() will use, based on the input angle. The output from the function *simple.rotation*() is a 3D array with the two joined data sets (data.1 and data.2). Please note that the two datasets are joined based on their associated specimen names (dimnames). Thus, the order of the specimens in each dataset is not important, as long as all the cases match perfectly between the two datasets in the same specimen. If there are extra specimens for one of the datasets or the names do not match properly, *simple.rotation*() will not include them in the output rotated joined dataset.

As explained in the previous section, the function used for the rotation step for example B needs four landmarks for each structure instead of three, as it is a more complex articulation that will need to get landmarks from both structure joined (or very close) to form the double-articulation hinges, as well as internally running separate rotations per structure on top of the final rotation. We will use *double.rotation*() function. For this function we will need to match the dimnames:

matched = match.datasets(data.3_t, data.4_t)

The datasets that will need to be rotated are the following:

data.3_rotate = matched$matched1

data.4_rotate = matched$matched2

Similarly to *simple.rotation()*, the *double.rotation()* will be as follows:

dr45l = double.rotation(data.1= data.3_rotate, data.2= data.4_rotate, land.a=1, land.b=2, land.c=3, land.d=4, land.e=1, land.f=2, land.g=3, land.h=4, angle=45) joined_dataset <-double.rotation(data.1 = translated_skull, data.2 = translated_mandible, land.a = 1, land.b=2, land.c=3, land.d=4, land.e=1, land.f=2, land.g=3, land.h=4, angle = 45)

The function double.rotation() will return a list that will need to be joined as it follows:

dr45 = join.arrays (dr45l$rotated1, dr45l$rotated2)

Double.rotation() allows the user to translate one of the objects after the rotation, in the case of not wanting them in contact to one another. For example:

skull_translate = c(1.7,0.1, 0)

dr45_st = join.arrays (dr45l$rotated1, translate(dr45l$rotated2,land.e, skull_translate))

### Plotting

Finally, after the rotation process we can visualize a 3D plot for each specimen with the two rotated structures for both a single-point and a double point articulations using the function *plot.rotation.3D()*:

plot.rotation.3D(joined.data= joined_dataset, data.1=data.1, data.2=data.2, specimen.num =1) plot.rotation.3D(joined.data=dr45_st, data.1=data.3, data.2=data.4, specimen.num =1)

The default plotting colours for the landmarks of the joined rotated structure (joined.data) is black and red for each original structure (data.1 and data.2).

### Exporting

After the rotation process we could either use the joined GM array in further analyses, visualise the resulting joined structure through *geomorph* (Adams & Otárola-Castillo 2013), or we could also export it and save it in order to use it in another software, such as MorphoJ (Klingenberg 2011). In this

step we will be using the function writeland.tps() in the R package *geomorph* (Adams & Otárola-Castillo 2013) in order to save a tps file from the joined GM array:

writeland.tps(A=“joined_arm”, file = “joined_arm.tps”, scale = NULL)

## Other applications

Our method is an important addition to the tool kit of the geometric morphometrics field. It will facilitate the analyses of compound 3D morphological datasets in geometric morphometrics analyses but will also be useful outside of this field as it can be applied to any method that uses 3D coordinates. The examples of applications are numerous in different fields of study, such as biology, anthropology, palaeontology, medical sciences, archaeology, and engineering. For example, in evolutionary biology, ShapeRotator would allow analyses of multiple or articulated hard structures (such as different segments of an exoskeleton, different articulated bones, or neighbouring plant structures, among others), different structures from the same object or organism (e.g. different and not adjacent body parts), or pieces from damaged specimens. In medicine and veterinary science it could be used to examine shape and size variation in different organisms’ growth due to different nutritional treatments or to examine how different structures respond to injuries or surgery. It would be useful in palaeontology or archaeology when trying to quantify shape of different objects or organisms that might have been preserved in disarticulated pieces. Finally, we would like to include a word of caution on how the angle chosen between different sub-structures could produce different results. As common-sense would suggest, the angle chosen should always be within the range of angles possible for that complex object, but in some cases it might be necessary to perform the GM analysis on more than one angle to be sure that the results do not differ. In other cases it might even be better to analyses sub-structures separately.

## Acknowledgements

We are grateful to two anonymous reviewers, Tom Semple, Zoe Reynolds, and Amit Kumar, for their helpful comments on the manuscript. JSK thanks the Australian Research Council for ongoing support. LB was supported by the Knut and Alice Wallenberg foundation, KAW 2013.0322 postdoctoral program in Mathematics for researchers from outside Sweden, and from SPP2026 from the German Research Foundation (DFG). This was part of the PhD thesis of MVG.

